# Validity of dynamical analysis to characterize heart rate and oxygen consumption during effort tests

**DOI:** 10.1101/2020.02.29.971119

**Authors:** D. Mongin, C. Chabert, A. Uribe Caparros, A. Collado, E. Hermand, O. Hue, J. R. Alvero Cruz, D. S. Courvoisier

**Affiliations:** Quality of Care Unit, University Hospitals of Geneva, Geneva, Switzerland; ACTES laboratory, UPRES-EA 3596 UFR-STAPS, University of the French West Indies, Guadeloupe, France; Malaga University, Andalucía Tech., Department of Human physiology, histology, pathological anatomy and physical education, Malaga, Spain; AVAE Laboratory, EA 6310, University of the French West Indies, Guadeloupe, France

## Abstract

Performance is usually assessed by simple indices stemming from cardiac and respiratory data measured during graded exercise test. The goal of this study is to test the interest of using a dynamical analysis of these data. Therefore, two groups of 32 and 14 athletes from two different cohorts performed two different graded exercise testing before and after a period of training or deconditioning. Heart rate (HR) and oxygen consumption (VO_2_) were measured. The new dynamical indices were the value without effort, the characteristic time and the amplitude (gain) of the HR and VO_2_ response to the effort. The gain of HR was moderately to strongly associated with other performance indices, while the gain for VO_2_ increased with training and decreased with deconditioning with an effect size slightly higher than VO_2_ max. Dynamical analysis performed on the first 2/3 of the effort tests showed similar patterns than the analysis of the entire effort tests, which could be useful to assess individuals who cannot perform full effort tests. In conclusion, the dynamical analysis of HR and VO_2_ obtained during effort test, especially through the estimation of the gain, provides a good characterization of physical performance, robust to less stringent effort test conditions.

## INTRODUCTION

Characterization of Heart Rate (HR) and oxygen consumption (VO_2_) related to mechanical power (i.e., speed or power) during standardized graded exercise test (GET) is an unavoidable step in current athlete’s performances assessment ^1^. These two measurements are also classically used in the scientific field of sport studies as one of the main physiological output to characterize evolution of athlete’s performance over time ^2–4^.

Current analysis of these parameters is based on two radically different approaches. The first is the use of standard techniques, easily applicable and extensively used. The most common index to characterize the HR recovery is the Heart Resting Rate (HRR) ^5^, commonly defined as the difference between HR at the onset of recovery and HR one minute after. This characterization is known to be a good predictor of cardiac problems in medicine ^5^, and is an interesting indicator of physical condition and training ^6^. The maximum rate of HR increase (rHRI) is a recent indicator showing correlation with fatigue and training in various studies ^6^.

This first type of approaches to characterize HR dynamics suffer from two important drawbacks. First, these measurements mix the amplitude of the HR response to effort with its temporal shape. For instance, someone reaching a maximum heart rate of 190 beat/minute and decreasing to 100 beat/min in one minute will have the same HRR as another person reaching 150 beats/minute and decreasing to 60 beats/minute in one minute, although the HR dynamic is different. Secondly and more importantly, they use only a small fraction of the information contained in the entire effort test (e.g., for HRR, the heart rate at the end of exercise and the heart rate one minute later, so two minutes out of a test of 20 to 30 minutes).

Regarding standard analysis of respiratory parameters, the main indicators of athlete’s performing capacities are the maximal VO_2_ reached during the exercise, the maximal aerobic power or the maximal speed reached, and the values of power or speed at the two Ventilatory Thresholds (VTs), corresponding to the lactic apparition (VT1) and the accumulation (VT2) threshold ^7^. Although these VO_2_ parameters are currently considered among the best indexes of aerobic fitness evaluation ^8^, determining them requires a visual analysis of the data and make use of only a part of the gas consumption dynamics, discarding the majority of the information contained in the entire effort test.

The second approach, based on dynamical system modeling, could allow to more accurately characterize the HR or VO_2_ response during effort. Dynamical analysis based on differential equations is an active subject of research in the behavioral field since the seminal work of Boker ^9^ and has led to numerous studies in the field of psychology and to several methodological advances ^10^. As approach based on first order differential equation approach show potential ability to adjust HR measurement ^11^ and VO_2_ dynamics during variable effort loads ^12^, we propose to use a simple first order differential equation coupled with a mixed effect regression to quantify the link between the exercise load during effort test and the resulting HR or VO_2_ dynamics. Because dynamical models use all the information measured during the effort test, it may allow to accurately assess performance indices using non-maximal effort tests.

The aim of this study is to characterize the indices produced by the dynamical analysis of HR and VO_2_ for different effort test protocols. Focus will be set on the link between the new dynamical indices and their standard counterpart (construct validity), to their ability to detect performance change over two different context of training load (predictive validity, and sensitivity to change), and their need for the full tests or for only the first part of the test (non-maximal effort test). Longitudinal data measured for two groups of young athletes with two different protocols will be analyzed. One group should show a performance increase following a three months training period, and the second group should have a performance decrease after an off-season of 6 weeks.

## METHODS

### Subjects

To test the reliability of the dynamical analysis model, data were acquired in two different populations (Guadeloupe and Spanish athletes) subjected to two different profiles of exercise (step-by-step cycling and continuous intensity running increase) and physiological conditions (training and deconditioning)

Group 1 consists of 32 young athletes (19 males and 13 females; 15.1±1.5 year-old) of the Regional Physical and Sports Education Centre (CREPS) of French West Indies (Guadeloupe, France), belonging to a national division of fencing, or a regional division of sprint kayak and triathlon. GET was performed at the end of the off-competition season, and after 3 months of intense training (3-7 sessions/week). All athletes completed a medical screening questionnaire and gave written informed consent prior to the study. The study was approved by the CREPS Committee of Guadeloupe (Ministry of Youth and Sports), the University Ethics Committee and performed according to the Declaration of Helsinki.

Group 2 consists of 14 young (males, 15.4±0.8 year-old) amateur soccer players from Malaga (Spain), performing three weekly training sessions and one weekly competition. A first GET was performed at the end of the soccer season and a second 6 weeks after. All participants were warned to avoid any training activity during this time. All athletes gave written informed consent prior to the study, and the measurements have been used in a previous publication ^13^.

### Effort test measurement

Group 1 performed an incremental testing on an SRM Indoor Trainer electronic cycloergometer (Schoberer Rad Meßtechnik, Jülich, Germany) associated to a Metalyzer 3B gas analyzer system (CORTEX Biophysik GmbH, Leipzig, Germany). Cardiorespiratory parameters were recorded cycle-to-cycle during all the testing to obtain HR and VO_2_ all along the test session. The effort protocol used consisted of a 3 minutes rest phase, followed by a 3 min cycling period at 50 watts, followed by an incremental power testing of +15 Watts by minute until exhaustion. Measurements of VO_2_, HR and mechanical power during the last increment sustained by athletes were respectively considered as VO_2_ max, HR max and Maximal Aerobic Power (MAP). At the end of the test, measurements were prolonged during a 3 min period to record the physiological recovery of athletes.

Group 2 performed GET on a PowerJog J series treadmill connected to a CPX MedGraphics gas analyzer system (Medical Graphics, St Paul, MN, USA) with cycle-to-cycle measurements of respiratory parameters -including VO_2_, and HR-with a 12 lead ECG (Mortara). The stress test consisted of an 8-10 min warm up period of 5 km.h^−1^ followed by continuous 1km.h-^1^ by minute speed increase until the maximum effort was reached. Power developed during the effort test was calculated using the formula described by the American College of Sport Medicine (ACSM). The latter determines an approximate VO_2_ of runners ^14^ associated to the Hawley and Noakes equation that links oxygen consumption to mechanical power ^15^.

### Truncated effort tests

In order to test the robustness of the dynamical analysis, truncated effort tests were generated from the maximal effort test for both groups. It consisted in removing the measurements of the test for power (or speed) above 2/3 of the maximum power (or maximum speed) value. The recovery period was set as the recovery measurements of the full effort test with values below the maximum value reached during the truncated exercise. An example of truncated effort is presented in Fig1, for a VO_2_ measurement during an effort test of group 1.

**Fig1:**
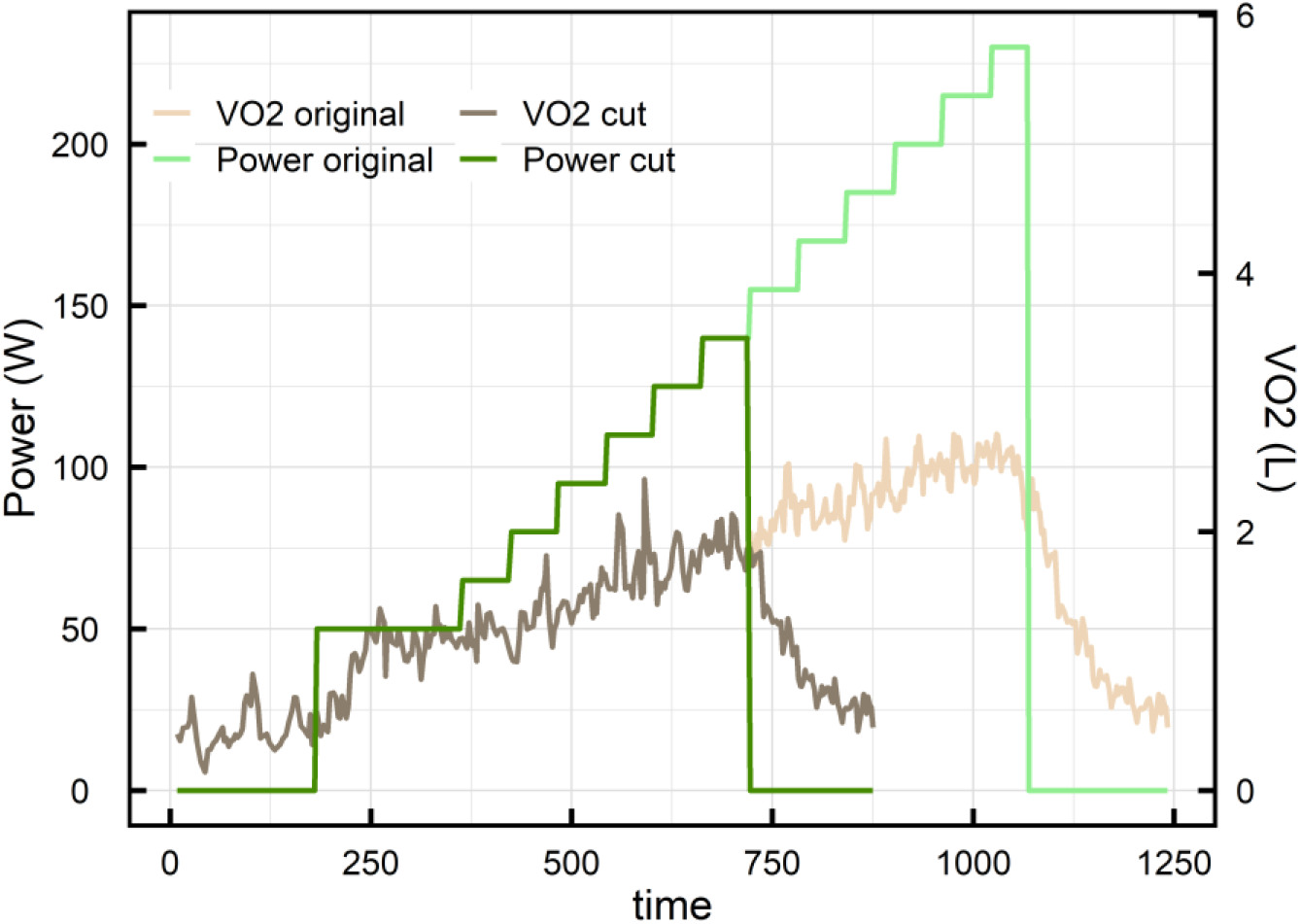
VO_2_ measured during a maximal effort test (light colors lines), and the truncated test generated from these data (dark color lines)

### Standard indices

The HRR calculated is the standard HRR60, which is the difference between the HR at the onset of the recovery and the HR 60 seconds later. The ventilatory thresholds 1 (VT1) and 2 (VT2) are calculated using the Wasserman method using the minute ventilation VE/VO_2_ for determining VT1 and VE/VCO_2_ for VT2 ^16^. The rHRI is derived by performing a sigmoidal regression of HR before and during the first 3 min effort step (only in group 1) and calculating the maximum derivative from the estimated parameters, as described in ^17^. Maximum aerobic power is the maximum power spend during the maximal effort test. HRmax and VO_2_ max are the maximum values of the rolling mean of HR and VO_2_ over 5 points.

### New indices using dynamical analysis

A first order differential equation describes a relation between the change in time of a variable, the value of this same variable and a possible time dependent excitation mechanism. For a variable *Y* (HR or VO_2_), it reads:

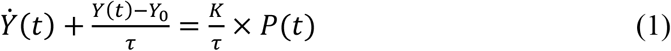

Where *Ẏ*(*t*) is the time derivative of *Y* (i.e. its instantaneous change over time), *Y*_0_ its equilibrium value (i.e. its value in the absence of any exterior perturbation) and *P*(*t*) the excitation variable, that is the time dependent variable accounting for the exogenous input setting the system out of equilibrium. Equation 1 describes the dynamics of a self-regulated system that has a typical exponential response of characteristic time *τ* and an equilibrium value *Y*_0_ in the absence of excitation (i.e. when *P*(*t*) = 0). For a constant excitation (i.e. a constant *P*(*t*) = *P*), the system stabilizes at a value *KP* after several *τ*. This value depends on both the system and the excitation amplitude (see Fig2 left panel).

**Fig2:**
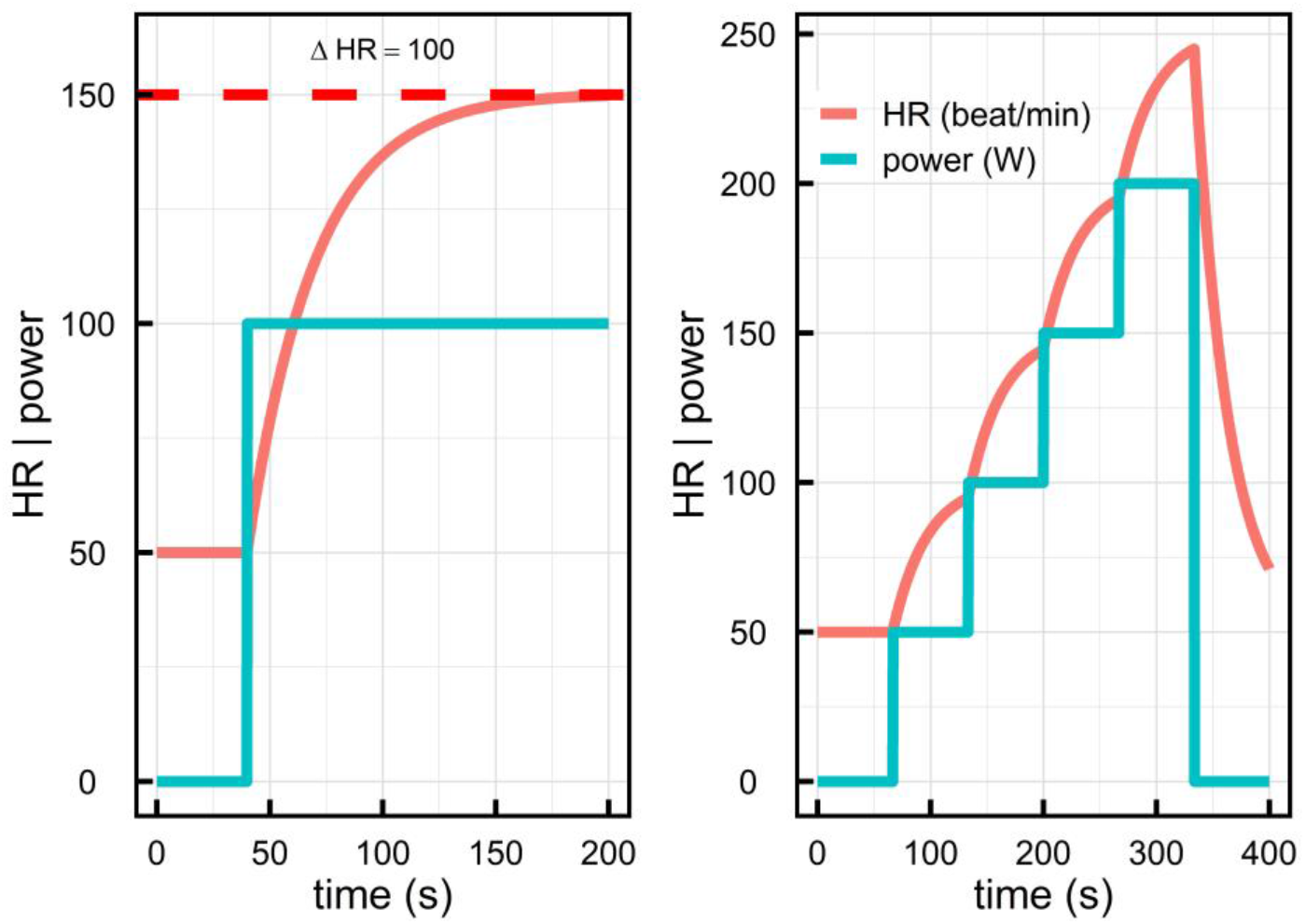
simulated HR dynamics following equation 1, for two different efforts (left panel: constant effort, right panel: effort test of three incremental steps), an equilibrium value of 50 beats.min^−1^, a decay time of 30 s and a gain of 1.

HR and VO_2_ are two self-regulated features of our body: they respond to an effort with a certain characteristic time to reach a value corresponding to the energy demand ^18^. Equation 1, as already demonstrated in ^12^ for VO_2_, can reproduce the dynamics of these two measures when considering that *P*(*t*) is the power developed by the body during effort. Assuming that HR or VO_2_ follow equation 1, only three time independent parameters are needed to characterize and to predict their dynamics for any time dependent effort:

- *Y*_0_ (i.e. *HR*_0_ or *VO*_20_) is the *equilibrium value*, i.e. the value in the absence of effort.
- *τ* is the characteristic time or *decay time* of the evolution of the variable. It corresponds to the time needed to reach 63% of the absolute change of value for a constant excitation. For instance, for an individual running at 10 km/h and who would have a total increase of HR of 60 beats/min for that effort, the decay time would be the time needed to increase his heartbeat by 38 beats/min (60 beats/min × 63%).
- *K*, the *gain*, is the proportionality coefficient between a given effort increase and the corresponding total HR or VO_2_ increase (Δ*HR* and Δ*VO*_2_). An illustration is provided in Fig2 left panel: a HR gain of *K*_*HR*_ = 1 beat/min/W leads to a HR increase of 100 beats/minute for a 100W effort increase, and to Δ*HR* = 200 beats/min for a 200W effort increase.

An example of the dynamics for HR following equation 1 is given in Fig2 considering *HR*_0_ = 50 beats.min^−1^, *τ*_*HR*_ = 30 s, *K*_*HR*_ = 1 and two efforts types. These three coefficients tightly characterize the dynamics of HR and allow us to generate the response to any effort.

The estimation of the three parameters characterizing the dynamics according to equation 1 is done in a two-step procedure, consisting in first estimating the first derivative of the variable studied over a given number of points with Functional Data Analysis (FDA) regression spline method ^10,19^. It consists on generating a B-spline function that fits the outcome to be studied and then estimating the derivative of that function. In order for the generated B-spline function to be differentiable, it needs to be smooth. This is achieved through a penalty function controlled by a smoothing parameter. This parameter was chosen to maximize the R^2^, which is the goodness of fit of the model to the data.

Once the derivative estimated, a multilevel regression is performed to estimate the linear relation between the derivative, the variable and the excitation. The above analysis procedure has been embedded and described in the open-source library *doremi* ^20^ available in the open source software R. Example code can be found in the example vignettes associated.

### Statistical analysis

HR measurements with a rate of change higher than 20 beat.min^−1^ from one measurement to the next one were first removed as spurious results from the sensors. Indices difference within each group between the first and the second measurement was assessed using paired t test, and effect sizes were estimated by Cohen’s d index. Associations between standard physical performance indices and the results of our dynamical analysis were assessed using Spearman rank correlation coefficients for continuous variables and logistic regression for dichotomous variables. Training was operationalized as a binary variable set to 0 for measurements before training for group 1 and after deconditioning for group 2 (untrained situation), and to 1 for measurements after training for group 1 and before deconditioning for group 2 (trained situation).

All analyses were performed using R version 3.4.2, the package doremi for the dynamical analysis and the packages *data.table*, *Hmisc* and *ggplot2* for the data management and statistical indicators.

## RESULTS

The associations between standard indices were high, especially between the maximum value of oxygen consumption (VO_2_ max), the MAP achieved and the ventilatory threshold powers for VO_2_ (correlations ranging from 0.73 to 0.93). There was also a significant negative correlation between rHRI and VO_2_ max (correlation coefficient of −0.42, p = 0.023), meaning that a higher maximum aerobic power reached during effort or a higher maximal VO_2_ is associated with a lower rate of HR increase during the first effort test (For full details of these associations, for the first time of measurement, see Supplementary Table 1 online).

Example of HR and VO_2_ dynamics is given in Fig3, together with the estimated curve obtained from the first order differential equation analysis. The model was very close to the observed values for both HR and VO_2_, and for both effort test protocols, with R^2^ (median [IQR]) of 0.96 [0.93, 0.97] for HR, 0.94 [0.92, 0.96] for VO_2_ in group 1, and 0.95 [0.91, 0.97] for HR, 0.94 [0.90, 0.96] for VO_2_ in group 2. The ensemble of the estimated values compared to the true observed ones are presented in Supplementary Fig1 (online).

**Fig3:**
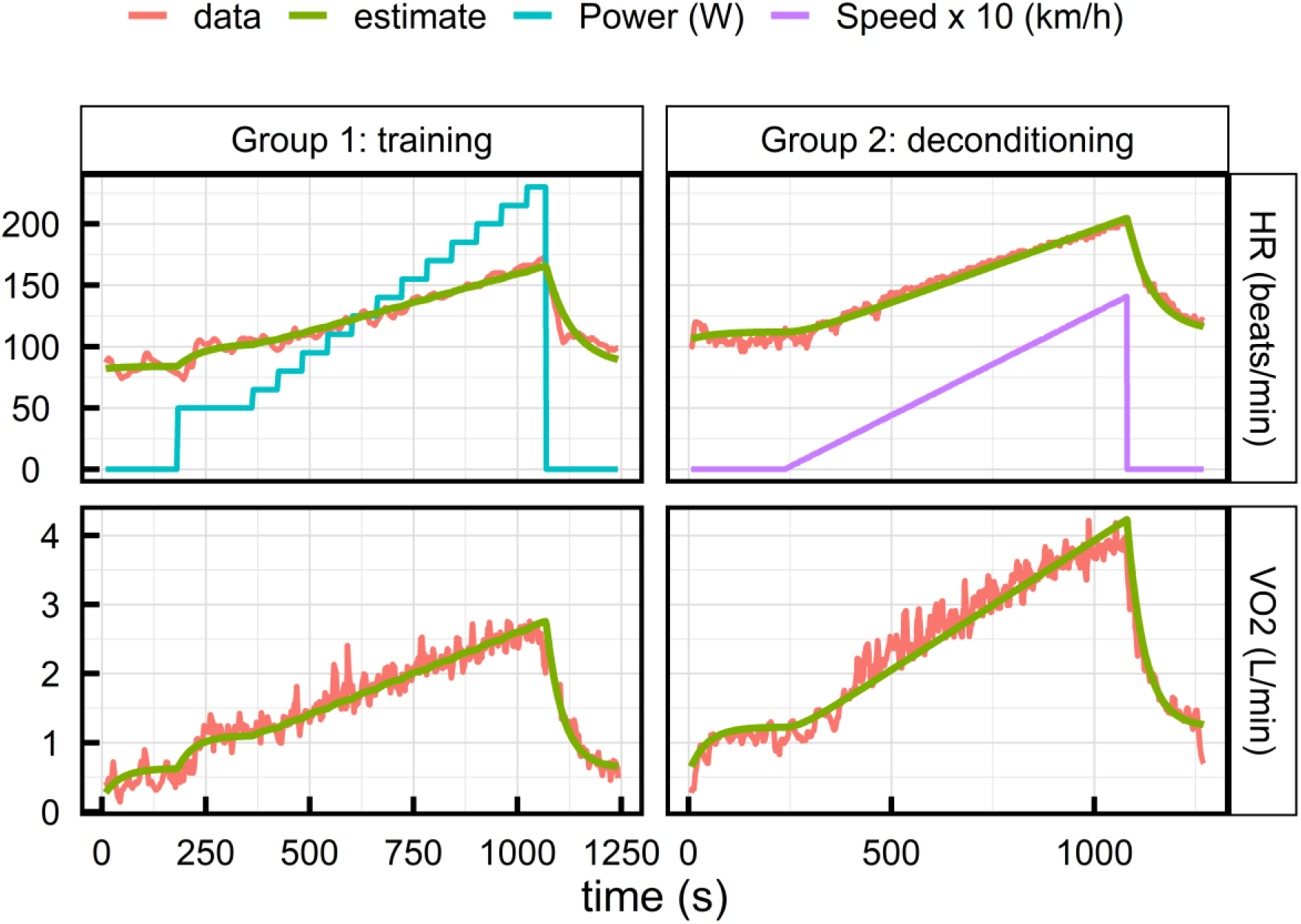
Example of HR and VO_2_ dynamics from one subject for each group. Blue line shows the power supplied by the subject during the effort, magenta line the speed on the treadmill, red lines are the experimental measurement of HR or VO_2_, and green lines shows the estimation given by the dynamical model.

The dynamical analysis estimation of resting values overestimated the measured values, certainly because the participants did not provide enough values before the start of the test (HR measures averaged approximately 20 seconds before the first effort increase, see Supplementary Table 2 online). Thus, we will discard this index for the rest of the study.

VO_2_ max and 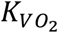, the gain of VO_2_ (i.e. proportionality coefficient between an effort increase and the final VO_2_ increase caused by this supplementary energy expenditure), increased significantly during the 3 months training period of group 1, and decreased significantly during the 6 weeks of deconditioning of group 2 (Table 1). The effect size was slightly higher for 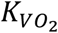 than VO_2_ max in the two groups and was higher for deconditioning than for training for both variables.

**Table 1.**
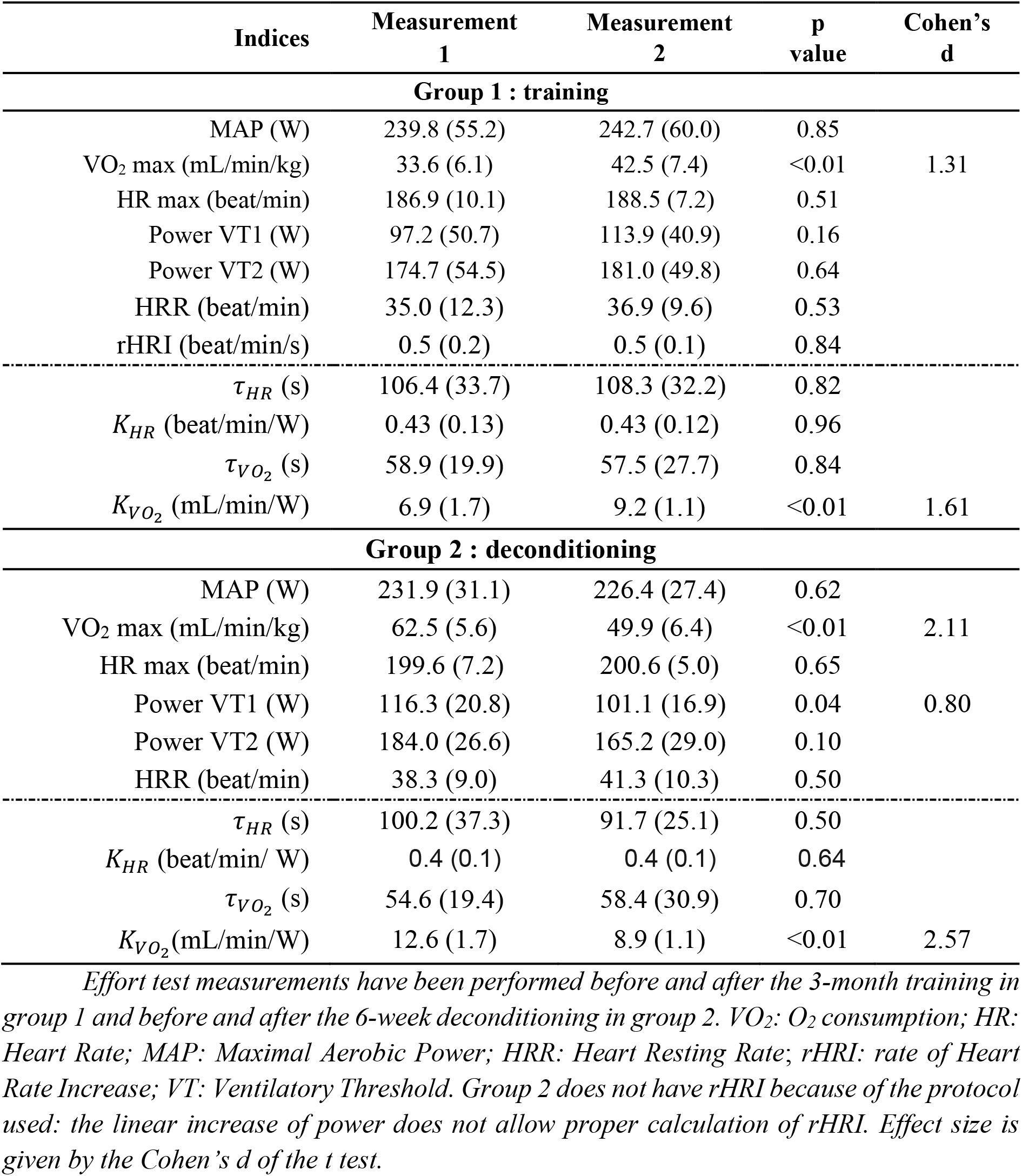
Comparison of the classical indices and the indices stemming from the dynamical analysis of VO_2_ and HR: the gain K and the decay time τ.

A small decrease of the power of the first ventilatory threshold (power VT1) is also observed in population 2. 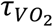, the response time *τ* of VO_2_ to the effort is shorter than *τ*_*HR*_, the response time of HR, in both populations. The HR gain (*K*_*HR*_) is remarkably similar in both groups, and unaffected by training or detraining.

In univariable analysis, *τ*_*HR*_ was correlated with measures of HRmax and HRR (Table 2), and *K*_*HR*_ (i.e., proportionality coefficient between effort increase and final HR increase caused by this supplementary energy expenditure) was negatively correlated with weight, maximal aerobic power, maximum O_2_ consumption, and the two ventilatory thresholds. Only in group 1, *K*_*HR*_ was also negatively correlated with age, height and rHRI, whereas a correlation with HRmax is found only in group 2. In other words, a decrease of *K*_*HR*_, (i.e. a decrease of Δ*HR* for a given effort) was linked with an improvement of oxygen maximal consumption, maximal aerobic power and the power corresponding at the two transition thresholds. Overall, correlations with new indices were higher than the correlations found between standard HR indices and other performance variables (see Supplementary Table 1 online). In a multivariable analysis performed in each group including age, weight, height, VO_2_ max and power at ventilatory thresholds, only weight remained significantly associated with *K*_*HR*_ (see Supplementary Table 3 online).

**Table 2.** Spearman correlation coefficients between the gain K and the decay time τ of HR and VO_2_ for both populations, physiological characteristics and standard analysis indices.

| Group 1 : training |  |  |  |  |
| --- | --- | --- | --- | --- |
| | $\tau_{HR}$ | $K_{HR}$ | $\tau_{VO_2}$ | $K_{VO_2}$ |
| Age | 0.11 | -0.43** | -0.01 | 0.06 |
| Weight | 0.07 | -0.73*** | -0.06 | 0.13 |
| Height | -0.06 | -0.55*** | -0.09 | 0.05 |
| MAP | 0.09 | -0.79*** | 0.12 | 0.01 |
| $VO_2$ max | 0.07 | -0.65*** | -0.10 | 0.57*** |
| Power VT1 | 0.13 | -0.44*** | 0.03 | 0.09 |
| Power VT2 | 0.10 | -0.63*** | -0.03 | 0.12 |
| HR max | 0.32* | 0.16 | 0.02 | 0.18 |
| HRR | -0.52*** | -0.14 | 0.06 | 0.06 |
| rHRI | -0.01 | 0.37** | 0.19 | -0.24 |
| Group 2 : deconditioning |  |  |  |  |
| Age | 0.07 | -0.08 | 0.10 | 0.20 |
| Weight | -0.11 | -0.63*** | -0.34 | 0.08 |
| Height | 0.16 | 0.05 | -0.16 | -0.13 |
| MAP | 0.28 | -0.73*** | -0.25 | -0.04 |
| $VO_2$ max | -0.05 | -0.49** | -0.25 | 0.67*** |
| Power VT1 | -0.05 | -0.54** | -0.15 | 0.19 |
| Power VT2 | 0.01 | -0.49* | 0.10 | 0.33 |
| HR max | 0.38* | 0.49** | 0.40* | -0.10 |
| HRR | -0.72*** | 0.16 | -0.32 | -0.08 |

MAP: Maximal Aerobic Power; HR: Heart Rate; HRR: Heart Resting Rate; VT: Ventilatory Threshold; rHRI: rate of Heart Rate Increase. Significance is indicated as follows: \*: $p < 0.05$ ; \*\*: $p < 0.01$ , \*\*\*: $p < 0.001$

VO_2_ decay time 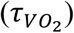 was globally independent of physiological variables and standard indices (Table 2), whereas 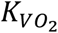 was strongly associated with VO_2_max. In a multivariable analysis performed on each group including age, weight, height, training, VO_2_max and power at ventilatory thresholds, VO_2_max and training remained significantly associated with 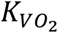 (see Supplementary Table 3 online). In group 1, training increased the 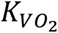 of 1.1 mL/min/W on average and an increment of 1L/min of VO_2_ max increased 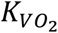 by 2.7 mL/min/W on average. In group 2, the deconditioning decreased 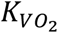 by 2.1 and the decrease of 1L of VO_2_ max lowered the VO_2_ gain by 1.8 mL/min/W.

### Truncated effort test

When performing the dynamical analysis on the truncated effort tests (see Fig1), the calculated R^2^ were slightly lower than the analysis performed on the entire test (0.90 [0.88, 0.94] for VO_2_ and 0.93 [0.89, 0.95] for HR in group 1, and 0.90 [0.87, 0.93] for VO_2_ and 0.90 [0.87, 0.95] for HR in group 2. The resulting dynamical indices were highly correlated with the one calculated on the entire effort test, as presented in Fig4.

**Fig4:**
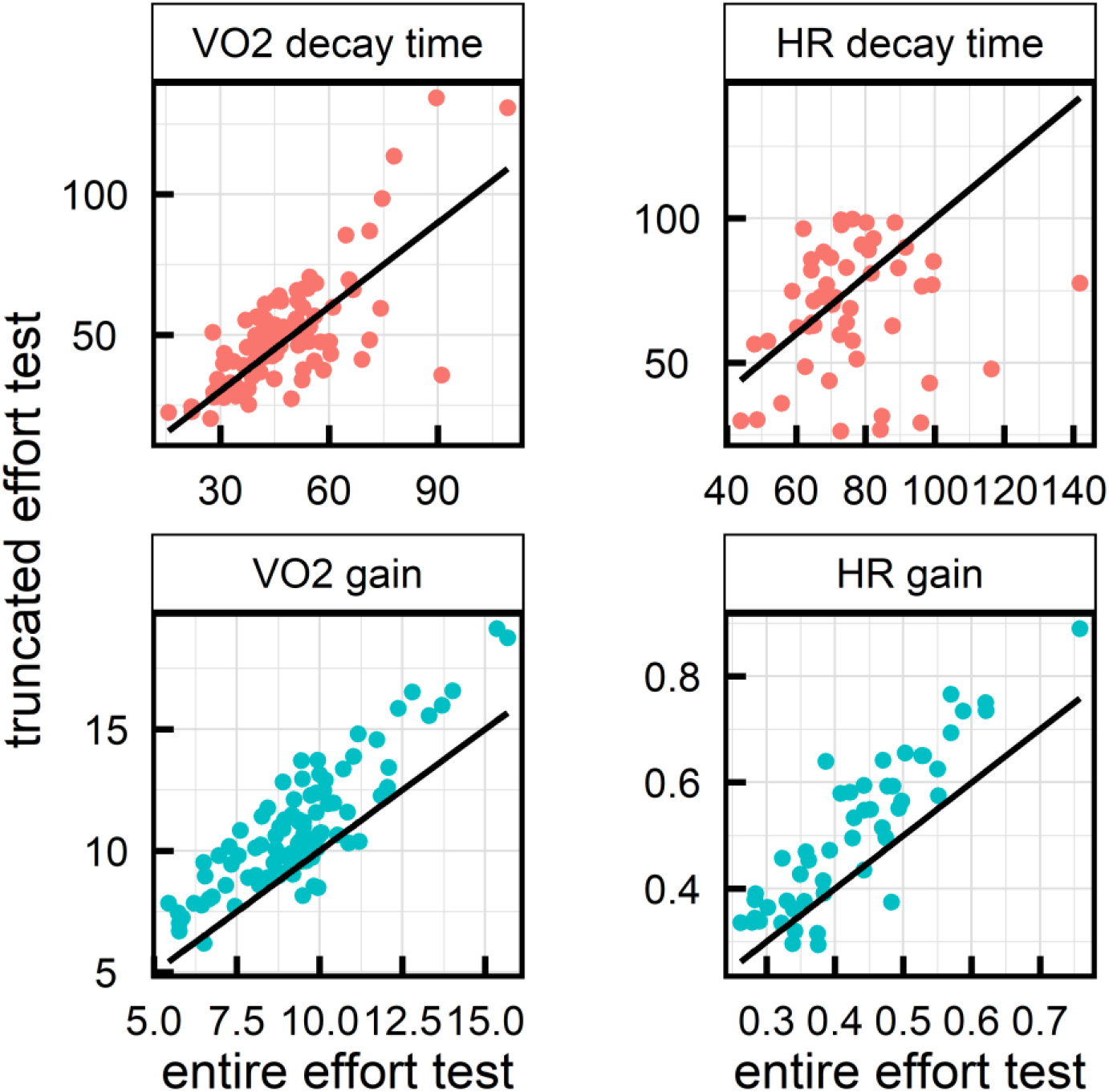
comparison of the dynamical indices estimated on the entire effort test (x axis) and on the truncated effort test (y axis) for VO_2_ (left column) and HR (right column). The solid black lines represent the identity.

The gain estimated on the truncated effort was slightly higher than the one estimated on the entire effort test. Correlation between the gain (for VO_2_ and HR) and the other performance indices remained similar to the ones observed in Table 2. The VO_2_ gain 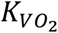 estimated on the truncated effort test still significantly changed between the two time points for both groups: from 8.9 (1.6) to 10.2 (1.8) mL/min/W for group 1 (p < 0.01, Cohen’s d = 0.754), and from 15.0 (2.3) to 12.2 (1.7) ML/min/W for group 2 (p<0.01, Cohen’s d = 1.38). In summary, the VO_2_ gain presented higher values but still significantly increased with training and decreased with deconditioning.

## DISCUSSION

### Main findings

Modeling the evolution of HR and VO_2_ during effort tests with a first order differential equation driven by the power spent during the effort produced an estimation able to reproduce at least 90% of the observed variance of HR or VO_2_. The model was successfully tested in two different populations (Guadeloupe and Spanish athletes) subjected to two different profiles of exercise (step-by-step cycling and continuous intensity running increase) and physiological conditions (training and deconditioning). The dynamical analysis provided three indices: the equilibrium value or resting state, the decay time, and the gain or proportionality between a given effort increase and the corresponding total increase in HR. HR gain was correlated to the main indices of athlete’s performance (MAP, VO_2_ max, VT1 and VT2), which was not the case of other standard HR indices. Furthermore, VO_2_ gain was sensible to change in training or physical deconditioning. Finally, the indices obtained when modeling truncated effort test (using about the first 2/3 of the effort tests data) had similar characteristics, showing the robustness and usefulness of such approach to incomplete effort tests. Such incomplete tests could occur due to lack of time but also when assessing older or sick persons.

### Standard indices

Using standard indices, it was possible to assess the relevance of the training/deconditioning conditions used for this study. Results were in line with those obtained by other studies ^6,18^, thus confirming the quality of the effort tests results in the two groups of athletes. In particular, the relationships between ventilatory thresholds (VT), maximal aerobic power (MAP) and maximum oxygen consumption (VO_2_ max), as well as the change in VO_2_ max after 3 months of training and after 6 weeks of deconditioning, were in accordance with expected results ^21^. VO_2_ max variation was also more pronounced in the deconditioning group than in the training one, as reported in previous observations ^22,23^. Concerning rHRI, the negative correlations with VO_2_ max and MAP was reported previously and is due to a parasympathetic withdrawal with sympathetic activation causing a relatively slower HR increase in response to intensity increase for well-trained athletes when compared to untrained ^3,6^.

### Dynamical analysis

As expected, there was a moderate correlation between VO_2_ gain 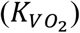 and VO_2_ max ^24^. Under an assumption of linearity between mechanical workload and O_2_ consumption, VO_2_ max corresponds to the oxygen consumption for the MAP expenditure and is directly linked to 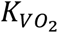:

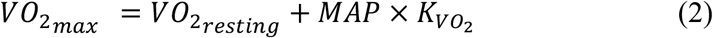

However, VO_2_ max is estimated via a single experimental measurement, supposed to be the VO_2_ at the maximum effort achieved by the athletes. The ability to reach maximum capacities during effort test is subject to several internal and external factors such as athlete’s engagement, mood state, fatigue and many others. Furthermore, the linear relation between energy demand and O_2_ consumption may not hold for high power expenditure ^25^, and thus VO_2_ max may not be representative of physical performance for intermediate efforts. In contrast, 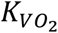 is estimated from the entire VO_2_ dynamics during the effort test, yielding a robust estimate of the VO_2_ response to effort, as shown by the fact that the VO_2_ gain estimated on truncated effort tests was still sensible to training and deconditioning.

The typical response time (i.e. the decay time *τ*) of VO_2_ was shorter than the HR one, in agreement with previous results ^26^. This temporal delay of HR compared to VO_2_ kinetics is due to the fact that heart flow regulation is partially driven by the oxygen demand of the organism detected via chemoreceptors, causing the HR increase to be a consequence of the VO_2_ increase ^27^.

The negative link between the *K*_*HR*_ and subject weight may be explained by the known association between fat-free mass weight and heart’s left ventricular size and mass ^28^. This association, reflecting a well-trained heart in heavier athletes, results in a lower ΔHR for a given effort and so a lower *K*_*HR*_.

### Strength and weakness

The main strength of this study is the use of two different populations of athletes, with two different effort tests and two different training schemes, showing its potential generalizability. Nevertheless, further study will need to extend these results to older adults, young children, and people with strong sedentary habits. A second strength is related to the analyses used, which allowed the estimation of performance indices without a maximum effort test. These analyses pave the way to obtaining accurate performance indices and information on training or deconditioning among larger groups of the population, such as the elderly, or patients at risk of cardiovascular events. The availability of ready to use, open source, tools for such analysis should facilitate its use for researchers and sport coaches ^20^.

As for limitations, the dynamic model used in this study made two assumptions that led to slightly suboptimal fits. First, the assumption that the equilibrium value is constant before and after the effort does not hold and led to the overestimation of these value. Indeed, HR and VO_2_ are known to decrease back to their resting value on a longer time scale due to the reduction of blood volume (i.e. dehydration), the evacuation of the heat accumulated during the muscular contractions, or the over-activation of the sympathetic system during exercise ^29^. The second assumption is that the entire dynamics has one unique characteristic exponential time, making the model unable to account for cardiac drift associated to prolonged effort or any long-term modification of the variable dynamics. However, despite the fact that the model can still be improved, it already provides indices with good sensibility to performance change and cardio-respiratory indices used to measure fitness.

## CONCLUSION

The dynamical analysis of heart rate (HR) and oxygen consumption (VO_2_) during effort appears to be relevant to evaluate performing capacities of athletes and its evolution. It reproduced at least 90% of HR or VO_2_ dynamics using only three estimated cardiovascular indices. It was more sensitive to training and deconditioning than classic indices. Furthermore, its ability to extrapolate VO_2_ and HR indices from truncated effort tests using only the first steps of the exercise could place it as a valuable tool for evaluate functional capacity from participants unwilling or unable to do maximal exercise testing.

