## Supplementary for "Validity of dynamical analysis to characterize heart rate and oxygen consumption during effort tests"

**Supplementary Table 1** Spearman correlation coefficients among standard performance indices. HR: Heart Rate; MAP: Maximal Aerobic Power; HRR: Heart Resting Rate; rHRI: rate of Heart Rate Increase (rHRI); VT: Ventilatory Threshold. Group 2 does not have rHRI because of the protocol used: the linear increase of power does not allow proper calculation of rHRI. Significance: *: p<0.5; **: p<0.001; ***: p<0.001.

| | **VO2 max** | **HR max** | **MAP** | **Power VT1** | **Power VT2** | **HRR** |
|---:|---:|---:|---:|---:|---:|---:|
| Group 1 : training | | | | | | |
| HR max | 0.16 | | | | | |
| MAP | 0.92*** | 0.20 | | | | |
| Power VT1 | 0.79*** | 0.35 | 0.81*** | | | |
| Power VT2 | 0.88*** | 0.26 | 0.93*** | 0.84*** | | |
| HRR | 0.11 | -0.03 | 0.08 | 0.19 | 0.12 | |
| rHRI | -0.42* | -0.12 | -0.36 | -0.21 | -0.36 | 0.18 |
| Group 2 : deconditioning | | | | | | |
| HR max | -0.03 | | | | | |
| MAP | 0.73** | 0.08 | | | | |
| Power VT1 | 0.75** | 0.21 | 0.73** | | | |
| Power VT2 | 0.88*** | 0.28 | 0.80** | 0.54 | | |
| HRR | 0.54 | 0.00 | 0.20 | 0.15 | 0.50 | |

**Supplementary Fig1** representation of the ensemble of the values estimated by the dynamical analysis compared to the true data, for HR and VO2 and the two groups. The red line is the identity function.

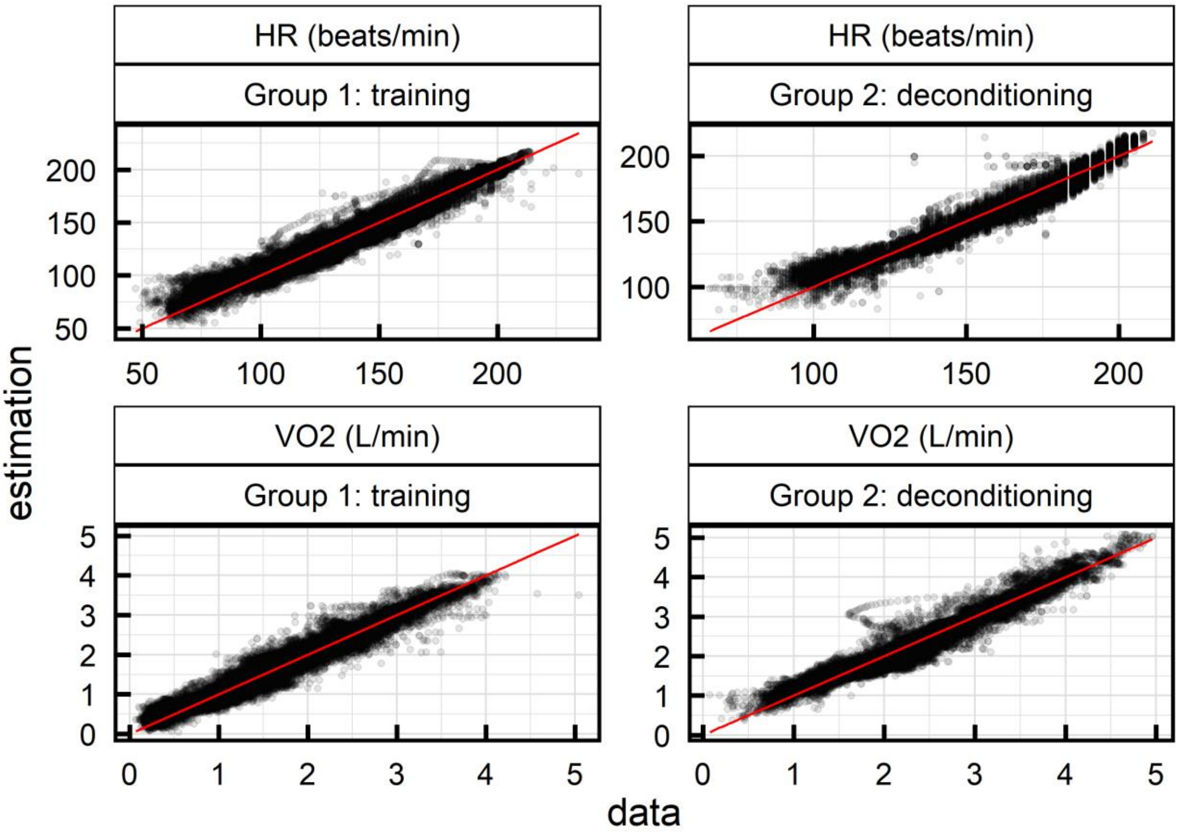

**Supplementary Table 2** Comparison between the estimated equilibrium value of HR ($HR_0$) and VO2 ($VO2_0$) and the mean initial value calculated from the experimental data before the test. Values reported are the mean variable values (sd).

| **variable** | First measurement | Second measurement |
|---|---|---|
| **Group 1 : training** | | |
| VO2 mean initial value (mL/min) | 366.1 (58.8) | 361.2 (68.8) |
| $VO2_0$ (mL/min) | 582.6 (116.6) | 585.5 (147.2) |
| HR mean initial value (beat/min) | 77.6 (10.0) | 81.7 (9.6) |
| $HR_0$ (beat/min) | 104.3 (9.6 | 104.7 (10.1) |
| **Group 2 : deconditioning** | | |
| VO2 mean initial value (mL/min) | 942.8 (87.5) | 1117.1 (140.1) |
| $VO2_0$ (mL/min) | 1225.8 (116.5) | 1384.0 (208.1) |
| HR mean initial value (beat/min) | 106.4 (5.6) | 103.6 (7.9) |
| $HR_0$ (beat/min) | 112.3 (7.4) | 111.1 (8.4) |

**Supplementary Table 3**: *multivariable analysis of the HR and VO2 gain for both groups. Training variable is set to 0 for the measurement taken in the untrained situation, and 1 for measurements taken in the trained situation. VT: ventilatory transition. Values reported are the estimated slope (standard error). Significance is indicated as follows: *: p<0.05; **: p < 0.01, ***: p < 0.001*

| | **HR gain (beat/min/W)** | | **VO2 gain (mL/W)** | |
|---|---|---|---|---|
| | **Group 1 training** | **Group 2 deconditioning** | **Group 1 training** | **Group 2 deconditioning** |
| **Age (year)** | -0.0024 (0.013) | -0.014 (0.014) | -0.028 (0.18) | 0.34 (0.29) |
| **Height (m)** | -9.3 (22) | 6.8 (29) | -1.6 (3.1) | -3.8 (5.4) |
| **Weight (kg)** | -0.0049* (0.002) | -0.0065* (0.0023) | -0.040 (0.032) | -0.012 (0.052) |
| **Training** | | | 1.1** (0.41) | 1.9** (0.63) |
| **VO2max (L/min)** | -0.056 (0.037) | -0.018 (0.032) | 2.8*** (0.52) | 1.8** (0.62) |
| **Power VT1 (W)** | -0.0002 (0.0005) | -0.0013 (0.00085) | -0.009 (0.0069) | -0.015 (0.016) |
| **Power VT2 (W)** | -3.9e-05 (0.00053) | -6.7e-05 (0.00068) | -0.012 (0.0074) | -0.0048 (0.013) |
